# Optimization of velocity receptor transduction in CAR T cells

**DOI:** 10.1101/2025.10.05.680548

**Authors:** Xinyun Jiang, Vasco Queiroga, Eban A. Hanna, Denis Wirtz

## Abstract

Limited infiltration capacity significantly limits the effectiveness of chimeric antigen receptor (CAR) T cells for solid tumors. We have recently developed a large family of highly modular synthetic cytokine receptors termed velocity receptors (VRs), capable of binding key inflammatory cytokines, such as IL5, IL8, and TNFα, which drive CAR T cells into an elevated motility state. These new CAR T cells sense and amplify these autocrine secreted cytokines, thereby maintaining a self-propelled, high migratory state, facilitating penetration into dense tumor cores. In this study, we systematically evaluated key factors influencing VR transduction in order to improve their stable integration and expression. We established a dual-fluorescence reporter system to allow simultaneous monitoring of both VR and CAR constructs, and while evaluating modifications to the vector construct and generating standardized infectious unit (IFU) curves under various conditions. Our results demonstrate that the attempt to reduce overall vector size by eliminating non coding sections upstream of the central polypurine tract (cPPT) do not yield better transduction efficiency, though it is unclear if the effect is due to viral production or integration impairment. We also observed a log-linear relationship between viral dose and transduction efficiency for a subset of VRs previously tested in various mouse models of human cancer, with VR5αIL8 and VR5αTNFα VRs consistently outperforming VR5αIL5 and V5 (full length native IL5 receptor). Overall, these findings establish an optimized and reproducible framework that offers valuable guidance for the future development and functional study of VR–CAR T cells in cellular therapies for solid tumors.

## INTRODUCTION

Chimeric antigen receptor (CAR) T therapy has demonstrated remarkable efficacy against hematological malignancies, including acute lymphoblastic leukemia^1^, non-Hodgkin lymphoma^2^, and multiple myeloma^3^. However, its efficacy in solid tumors remains limited due to immunosuppressive microenvironmental barriers, heterogeneous antigen expression, and insufficient CAR T-cell infiltration^4^. To overcome these challenges, our laboratory has engineered a class of synthetic cytokine receptors termed velocity receptors (VRs), which can harness self-secreted cytokine signals such as IL-5, TNFα, IFNγ, and IL-8 to lock CAR T cells into a high-motility state^5,6^, enabling them to penetrate dense stroma and infiltrate deeply into tumor cores^7,8^. This strategy significantly enhances the intrinsic migratory capacity and anti-tumor function of CAR T cells. The design of VRs - which contain binding, hinge, transmembrane, and signaling domains - is extremely modular^7^, allowing for more than 30k combinations of the various domains. We have made and tested *in vitro* and *in vivo* models of pancreatic cancer, ovarian cancer, and lung cancer using CAR T cell therapies transduced with a small subset of these constructs, including VR5αIL8, VR5αTNFα, VR5αIL5 and V5 (the full-length native receptor for IL-5) ^7^.

In the generation of CAR T cells, lentiviral (LV) vectors are commonly used to stably integrate exogenous genes into the T cell genome. As enveloped RNA viruses derived from the retroviridae family, LV vectors can efficiently enable long-term, stable expression introduced genetic material^9–11^. They are also the primary vector type currently approved by the FDA for CAR T-cell manufacturing^12,13^. This approach allows efficient gene transfer into both stationary and dividing cells and is well suited for sustained expression^14,15^, particularly with single-vector transductions with relatively small inserts^16^. However, introducing both VR and CAR constructs into the same cell requires dual LV transductions, involving two rounds of infection that increase experimental complexity. This often results in heterogeneous positivity within the cell population^17^ and makes it difficult to monitor the expression of both vectors simultaneously.

In particular, VR constructs often display reduced transduction efficiency and poor consistency across donors. To address these challenges, this study systematically optimizes VR transduction in T cells. We first introduced a GFP reporter downstream of the VR to enable synchronous and real-time monitoring of VR and CAR expression. Subsequently, we tested truncated VR constructs and modified the placement of the central polypurine tract (cPPT), a pivotal regulatory element, to investigate how these changes affect transduction efficiency and viral yield. Finally, we assessed different donor sources, detection strategies (antibody versus GFP reporter), and culture conditions to generate infectious unit (IFU) curves, thereby establishing a standardized framework to guide future VR design and application (**Fig. 1**).

**Figure 1.**
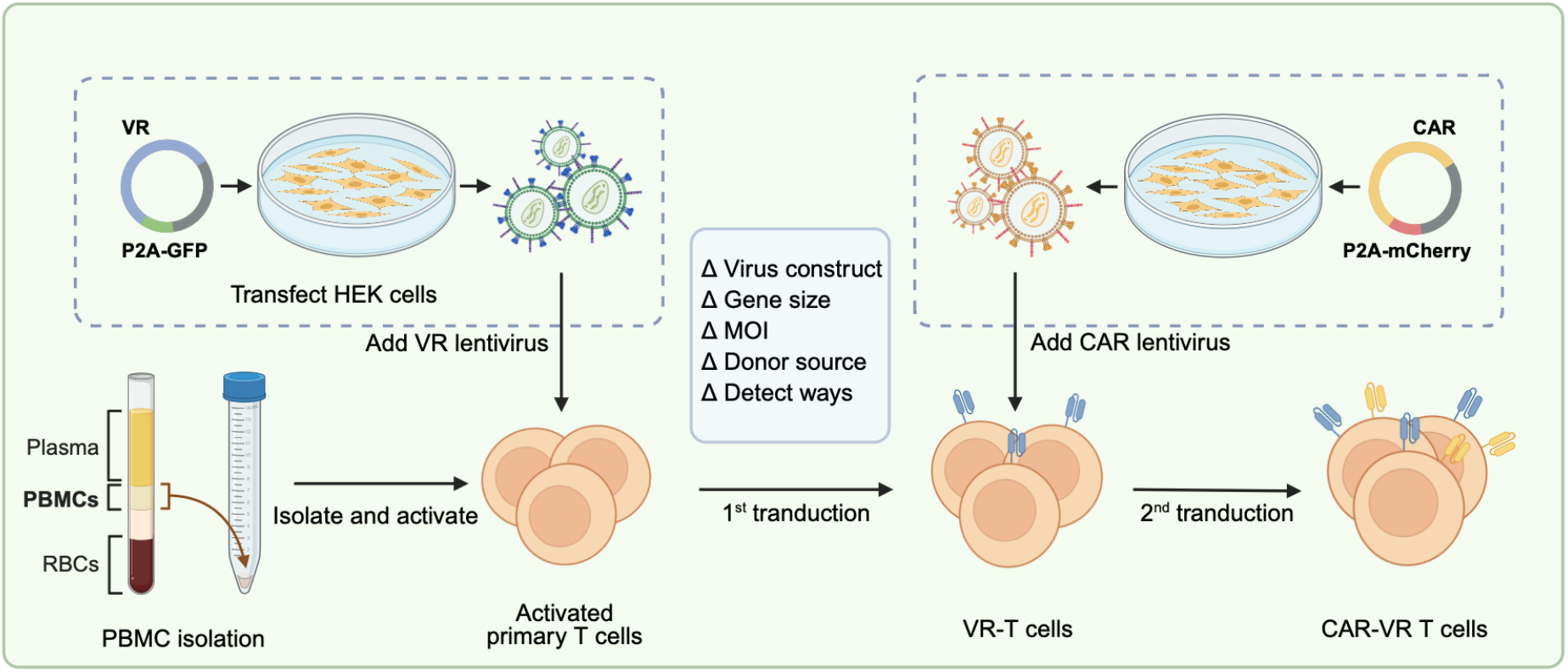
Schematic overview of VR-CAR T cell transduction. Plasmids are transfected into HEK293T cells to generate lentiviral particles, which are subsequently harvested and concentrated. Peripheral blood mononuclear cells (PBMCs) are isolated from leukopaks using Ficoll density gradient centrifugation. T cells are then isolated, activated, and sequentially transduced with VR and CAR vectors. After expansion, transduced T cells are assessed to confirm receptor expression. In the first VR transduction step, we primarily optimized viral construct size, multiplicity of infection (MOI), donor source, and detection methods.

## RESULTS

### Fluorescent labeling allows for real-time assessment of velocity receptors and CAR expression

In its initial implementation, detection of VR transduction efficiency relies on antibody staining^7^. However, due to the similarity of scFv regions within the VR and CAR extracellular domains (**Fig. 2**), it is difficult to select antibodies that discriminate between them. This results in cross-reactive staining of VR and CAR on the surface and confounded readouts. To avoid this issue, VR transduction efficiency is measured immediately after the first VR transduction, while GFP expression is used to assess CAR transduction after the secondary transduction, assuming that VR efficiency remains constant between the two round and for all subsequent experiments. Nevertheless, this assumption is approximate because the second transduction may alter cell state or interfere with stable VR expression, ultimately affecting transduction efficiency. Additional viral exposure can stimulate or damage T cells, leading to changes in cell viability and functional status while newly integrated genetic material can interfere with existing transgene expression^17,18^. Specifically, lentiviral pathogen-associated molecular patterns can engage pathways such as TLR7/8, RIG-I/MDA5, and cGAS–STING, placing T cells under stress. Such stress might further increase reactive oxygen species and activate cellular damage responses. Additionally, receptor expression on a cell’s surface can change over time even without antigen exposure^19^. Consequently, the difficulty in accurately determining the co-expression levels of both vectors within the same cell directly limits the reliability of downstream functional assessments of VR-CAR T cells.

**Figure 2.**
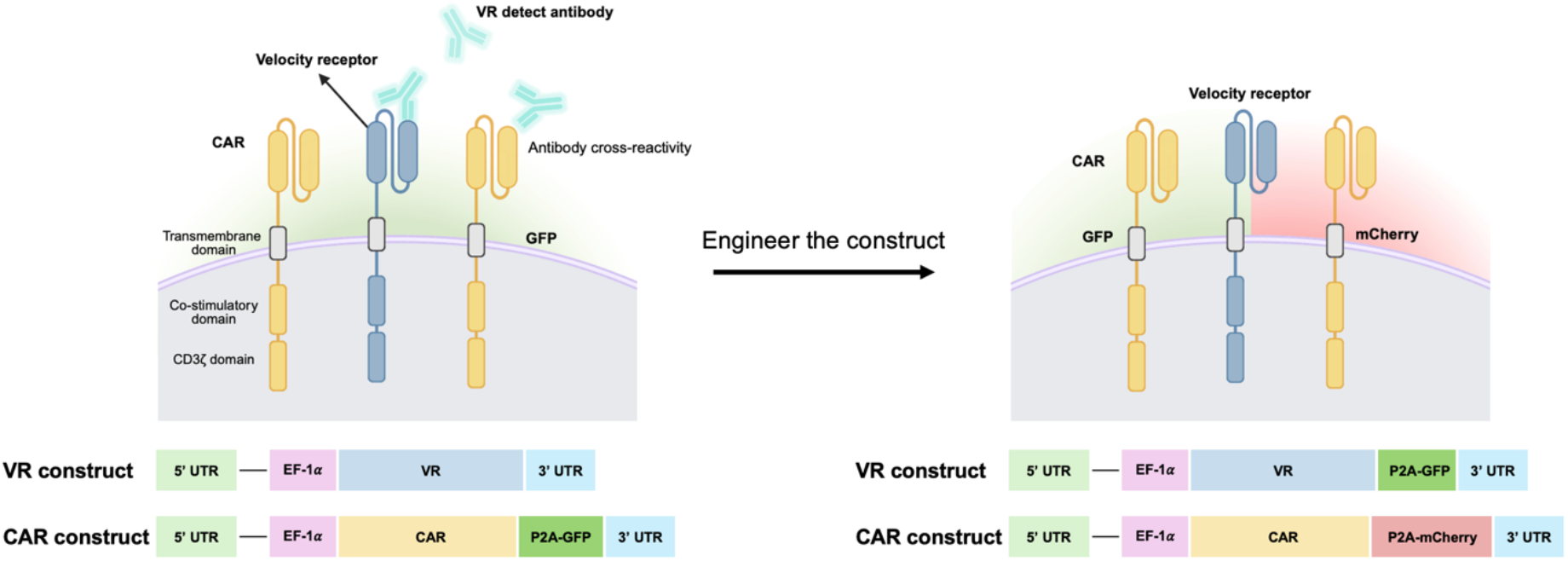
Dual-reporter system enables simultaneous detection of CAR and VR expression. Conventional antibody-based detection of VR expression can cross-react with CAR, resulting in inaccurate quantification of co-expression. To overcome this, GFP was incorporated into the VR vector via a P2A sequence, allowing direct fluorescence-based readout. This design enables simultaneous and real-time monitoring of VR and CAR expression in transduced T cells, while eliminating antibody cross-reactivity and simplifying flow cytometry analysis.

Therefore, we inserted a GFP reporter downstream of the VR sequence via a P2A fragment, enabling real-time quantification of VR transduction efficiency, and applied this strategy to all tested VR constructs. We employed a CAR vector carrying an mCherry reporter, in parallel, for T-cell transduction to eliminate spectral interference during detection (**Fig. 2**). This dual-reporter system not only allows simultaneous and real-time monitoring of VR and CAR expression, but also provides a more accurate basis for subsequent functional analyses while simplifying the flow cytometry workflow compared with antibody-based detection. Furthermore, fluorescent tags on both receptors allow for visualization and quantification of their expression *in vitro* during live imaging and *ex vivo* via cryosections of tumors or other organs.

### VR vector length influences viral titer and transduction efficiency

Previous studies have shown that LV transduction efficiency is strongly influenced by gene length, with longer constructs generally associated with reduced efficiency^20^. This effect reflects the size of the entire expression cassette, including the 5′LTR, coding sequence and 3′LTR, rather than the coding sequence alone^21^. In HIV-1–based LV vectors, the expression cassette usually contains a fragment of the *pol* gene located between the 5′LTR and the transgene. This fragment includes the cPPT, an element critical for nuclear import of the pre-integration complex. Although prior reports indicate that repositioning the cPPT can still preserve transduction efficiency, its local context remains poorly understood^22^.

Based on this, we attempted to delete non-essential sequences adjacent to the cPPT in order to enhance VR transduction. We generated truncated versions of the VR5αIL5 and V5 vectors (**Fig. 3A**), reducing their lengths from 7,872 bp to 7,039 bp for VR5αIL5 (a 10.6% reduction) and from 7,823 bp to 7,023 bp for V5 (a 10.2% reduction). Viral particles were subsequently produced in HEK293T cells in a single production batch and used to transduce T cells at varying volumes to assess dose-dependent transduction efficiency. Unexpectedly, under these conditions, V5-cut exhibited markedly lower efficiency compared with full-length V5, whereas VR5αIL5-cut achieved levels comparable to VR5αIL5 (**Supplementary Fig. 1**). RT-qPCR quantification further revealed that viral titers decreased by approximately 100-fold for V5-cut and 5-fold for VR5αIL5-cut compared to their parental constructs (**Fig. 3B**). Even after normalization to viral genome copy number, V5-cut transduction efficiency remained lower than that of V5 and VR5αIL5-cut showed a minimal improvement over VR5αIL5 (**Fig. 3C**).

**Figure 3.**
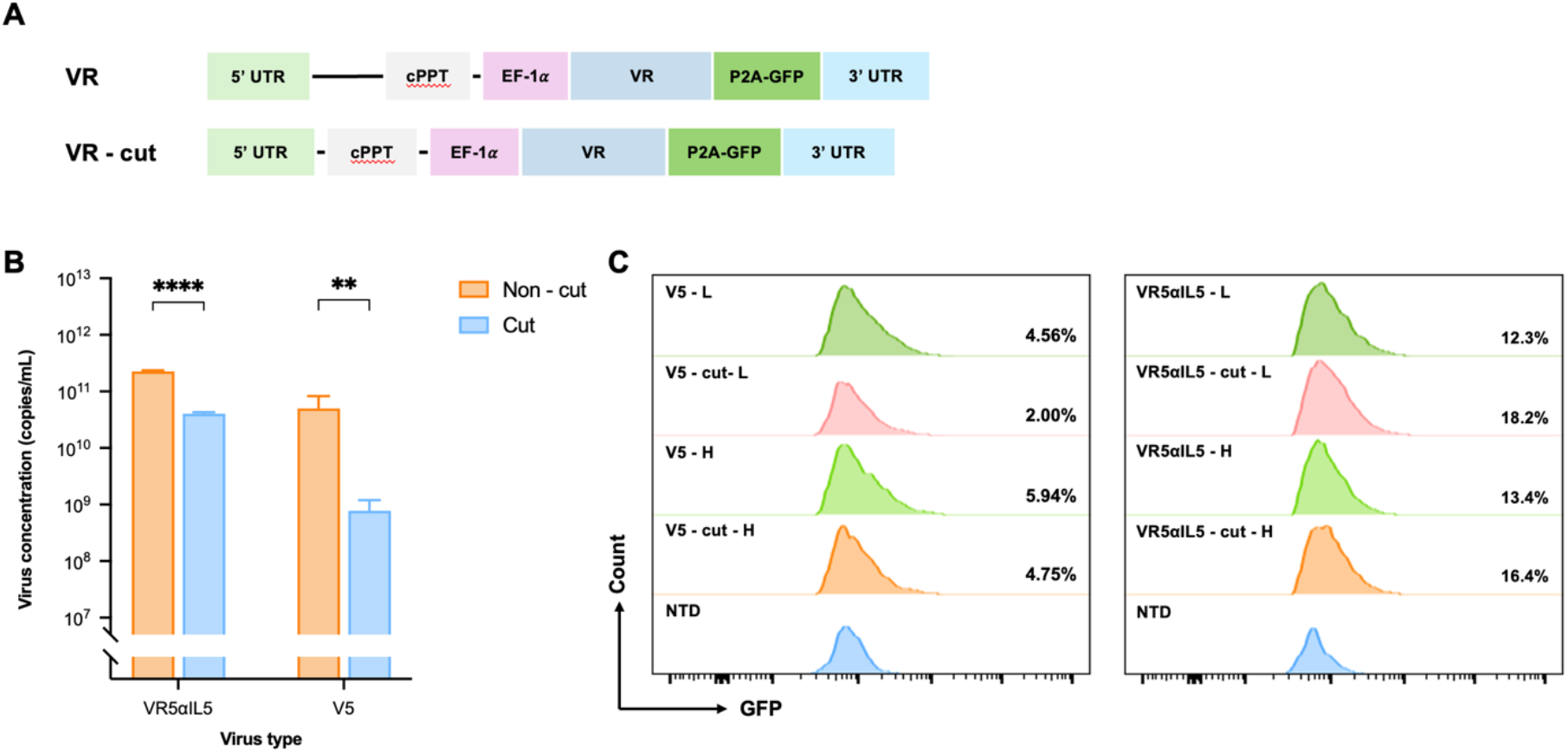
Comparison between truncation of VR constructs and parental on viral production and transduction efficiency. (A) Schematic of full-length VR and truncated VR (VR-cut) constructs, in which non-essential sequences adjacent to the cPPT were deleted. (B) Viral titers determined by RT–qPCR showed that truncation markedly reduced production, with V5-cut decreasing by ~100-fold and VR5αIL5-cut by ~5-fold compared to their parental constructs. Data are presented as mean ± s.d., and statistical significance was assessed using two-way ANOVA (****P < 0.0001, **P < 0.01). n = 4 technical replicates. (C) Flow cytometry histograms of GFP expression in T cells following transduction with full-length or truncated VR constructs at low (L) or high (H) viral doses at Day 6. NTD, non-transduced control.

These findings indicate that deletions near the cPPT, which further shift its position away from the vector center, severely restrict viral production and subsequent transduction. This suggests that maintaining the cPPT in a central position is essential, particularly in longer constructs such as V5. Considering both viral yield and transduction performance, we therefore used the unmodified parental vectors for subsequent optimization and functional studies.

### Optimized culture conditions establish reproducible IFU curves

To improve experimental reproducibility and ensure reliable results, we optimized transduction under multiple conditions and established IFU curves. T cells were isolated from PBMCs of different donors and separately transduced with four VR constructs (VR5αIL8, VR5αTNFα, VR5αIL5 and V5), with transduction efficiency measured on days 6, 12, and 18 to generate IFU curves. Transduction efficiency rose steeply at low doses and plateaued at higher doses, showing an approximately log-linear dose–response over the range. In both donors, transduction efficiency for all four VRs increased from day 6, peaked at day 12 and declined by day 18. Clear differences were also observed among VRs, with VR5αIL8 and VR5αTNFα consistently showing the highest efficiencies, VR5αIL5 intermediate, and V5 the lowest (<5%). Within the same experimental batch, no significant differences were observed between donor 6 and donor 7 at most time points, suggesting that donor variability had a relatively limited impact on VR transduction efficiency (**Fig. 4A, B**). Moreover, in a separate transduction experiment using the same viral batch, donor 8 exhibited higher efficiencies across all four VRs at day 18 compared to donor 6 and donor 7, indicating that batch-to-batch variability may exceed donor-to-donor variability (**Fig. 4C**).

**Figure 4.**
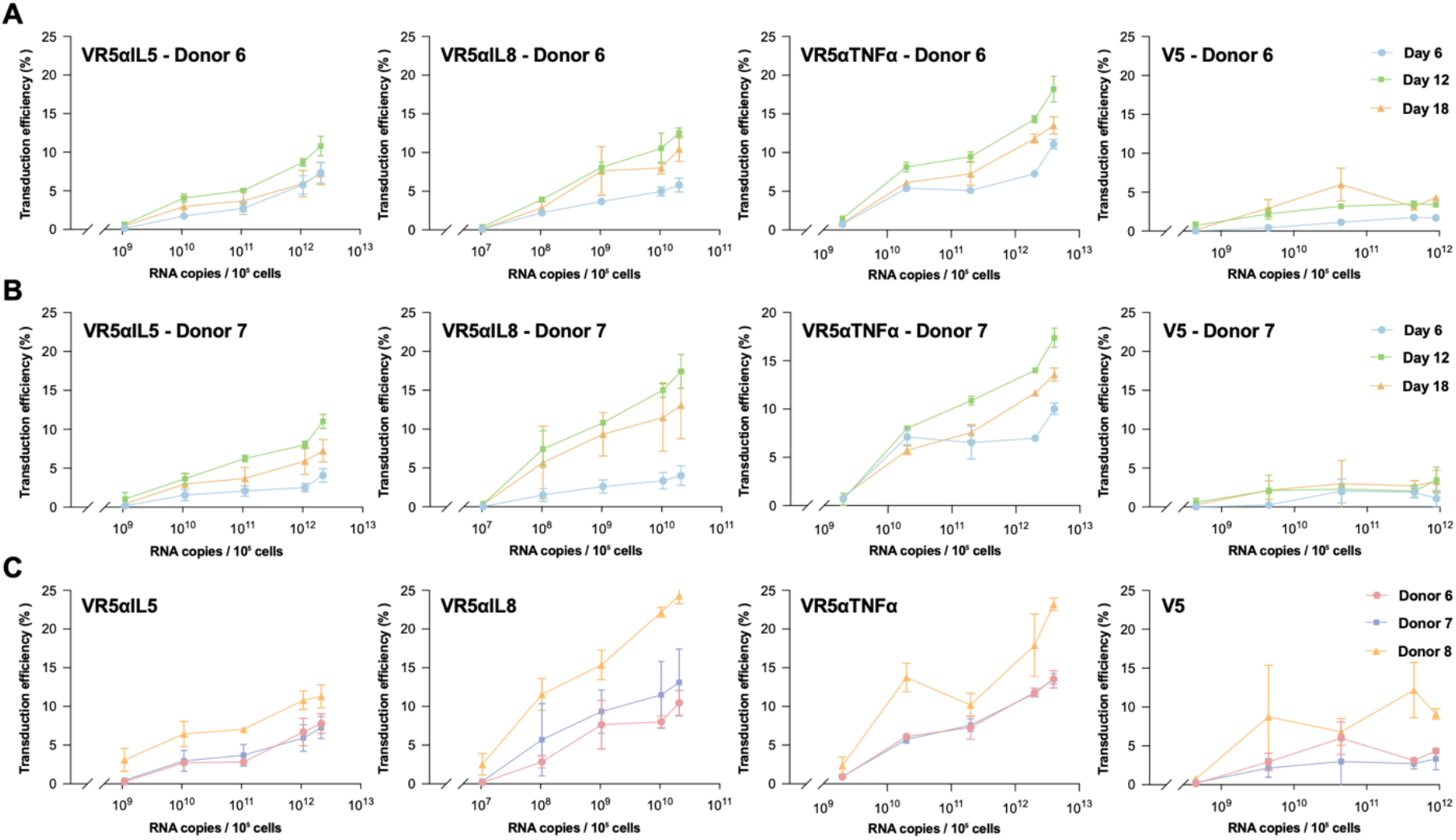
IFU curves of VR transduction efficiency across donors and time points. (A) T cells from donor 6 and donor 7 (B) were transduced with VR5αIL5, VR5αIL8, VR5αTNFα, or V5 vectors at different viral doses, and transduction efficiency was assessed on days 6, 12, and 18. (C) Comparison of transduction efficiency among donors 6, 7, and 8. Donors 6 and 7 were transduced in the same experiment, while donor 8 was transduced in a separate experiment using the same viral batch. Transduction efficiency was assessed on days 6, 12, and 18. Data represent mean ± s.d.. n = 3 technical replicates.

We further examined whether different spin-down speeds during viral loading on retronectin-coated plates would influence viral attachment and subsequent transduction efficiency. Higher centrifugation speed enhanced VR5αTNFα transduction efficiency at day 6, but resulted in a slight decrease at day 12, while V5 showed no significant changes under different conditions (**Supplementary Fig. 2**). Finally, we compared antibody-based and GFP-based detection of receptor expression and found that GFP proved to be a reliable measure of transduction efficiency and receptor expression with the expression being similar between the two measurements (**Supplementary Fig. 3**). This result confirms the reliability of fluorescent tags to assess as an antibody-free way of VR expression measurement.

## DISCUSSION

In this study, we established a dual-fluorescence reporter system by inserting GFP downstream of the VR sequence, enabling real-time monitoring of both VR and CAR expression and overcoming the limitations of antibody-based detection. Through linking GFP via a P2A sequence, the system provided a direct and stable assessment of post-transcriptional expression. Besides, we compared GFP-based and antibody-based detection of transduction efficiency and found GFP is reliable for detection. Some staining did not work well, which is likely because staining is influenced by many variables, including epitope access, washing, and timing. Using GFP as the transduction efficiency detection also simplifies cell preparation before flow cytometry.

Based on this system, we systematically evaluated key factors influencing viral yield and VR transduction efficiency. Previous studies have noted that LV packaging capacity is limited to approximately 8–9 kb, and oversized vectors can compromise packaging efficiency and titer^21^. While earlier work has emphasized the presence of cPPT and its role in nuclear import^23–25^, few studies have investigated how altering its position affects vector performance. Our results demonstrated that simply shortening sequences adjacent to the cPPT does not improve transduction efficiency but instead remarkably reduces viral titers. Thus, the relative position of the cPPT and its local sequence context play a critical role in determining viral yield and transduction efficiency, particularly evident in the bigger construct like V5. A possible explanation is that displacement of the cPPT from the central region or deletion of its flanking sequences disrupts structural integrity, thereby impairing nuclear import and reducing transduction efficiency. This observation implied that in VR optimization, vector architecture and regulatory element positioning are more critical than overall gene length reduction.

In addition, IFU curve analysis showed an approximately log-linear increase in transduction efficiency over the dose range, with a plateau at higher virus doses. This might because of the saturation of low-density lipoprotein receptor mediated entry sites on T cell surface^26^ and further competition from non-infectious empty viral particles^27^. VR5αIL8 and VR5αTNFα remained consistently the highest efficiencies and V5 the lowest. These findings establish IFU curves as a reliable quantitative reference for VR–CAR T studies, indicating broad applicability given the minimal donor-to-donor variability, though batch-to-batch variability remained more pronounced. We further found that minimizing cellular damage and loss during medium changes on the first day post-transduction, as well as changing the medium one day before the efficiency measurement, could help to improve transduction outcomes.

In summary, we enabled real-time detection of VR expression and systematically assessed how genetic structure, and experimental conditions influence viral production and transduction efficiency. This work establishes a more reliable and reproducible framework for VR-based T-cell engineering. Despite these findings, only certain relevant factors were examined here. Other parameters such as retronectin coating concentration, duration of T-cell activation^28^, viscous medium concentration^29^, and IL-2 concentration^30^ were not investigated in depth. Moreover, the generation of VR–CAR T cells typically require a second round of LV transduction, a process that may introduce additional variability, including the CAR to VR ratio, transduction order and potential vector competition, which warrants further study. A tandem gene construct could also be considered by splitting the bicistronic construct into two individual cassettes flanking the cPPT, thereby re-centering the cPPT and improving co-expression consistency while preserving nuclear import and transduction efficiency.

## ACKNOWLEDGEMENTS

This work was supported by grants from the National Institutes of Health (U54CA268083 and RO1CA300052, UG3CA275681) and the Patrick C. Walsh Prostate Cancer Research Fund at the Johns Hopkins School of Medicine. All cartoons were created with BioRender.com. Data analysis and graphing were performed in GraphPad Prism 10.

## Methods

### Cloning of P2A-GFP into VR vectors

VR nucleic acid sequences were synthesized as previously described in our earlier work. The P2A-GFP fragment was amplified from the laboratory’s M5 CAR-P2A-GFP construct. VR constructs were linearized by PCR with primers containing 15–20 bp homologous overlaps to the P2A-GFP insert. PCR products were confirmed by agarose gel electrophoresis and purified with QIAquick PCR Purification Kit (Qiagen, 28106). Infusion was performed using the In-Fusion Snap Assembly Kit (Takara, 638947). Purified VR backbones and P2A-GFP inserts were combined and incubated at 50 °C for 15 min. 2.5uL of the assembly mixtures were transformed into chemically competent E. coli (Invitrogen, C737303) and plated on LB agar supplemented with 100 µg/mL ampicillin (Quality Biological, 340-108-231). Following overnight incubation at 37 °C, individual colonies were picked for screening. Correct clones were identified by colony PCR with primers spanning the VR–P2A-GFP junction, followed by agarose gel electrophoresis. Correct clones were cultured in LB broth with 100 µg/mL ampicillin (Sigma-Aldrich, A5354), and plasmids were isolated using a QIAquick Spin Miniprep Kit (Qiagen, 27106). After sequencing, verified constructs were further amplified by QIAquick Spin Midiprep Kit (Qiagen, 28106) to obtain enough for applications.

### PBMC isolation

Peripheral blood mononuclear cells (PBMCs) were isolated from leukopaks sourced from healthy adult donors via the Blood Donation Center at Anne Arundel Medical Center (Annapolis, MD, USA), using Ficoll-Paque Plus (Cytiva, 17144003) according to the manufacturer’s instructions. For each preparation, each 5mL blood samples were mixed with 30 mL of DMEM (Corning, 10-013-CV), then gently added 13 mL of Ficoll-Paque Plus beneath the mixture, ensuring minimal disturbance at the interface. Samples were then centrifuged for 30 minutes at 400 g to achieve clear separation into plasma, PBMCs, Ficoll-Paque Plus, and erythrocyte layers. The layer containing PBMCs was carefully collected, pooled, and re-centrifuged at 400 g for 10 minutes. Cell counts were determined using 3% acetic acid with methylene blue staining solution (Stemcell, 07060). PBMCs were then adjusted to a final concentration of 5 × 10^7^ cells/mL in a cryopreservation solution consisting of 90% FBS (Corning, 35-010-CV) and 10% DMSO (ATCC, 4-X). Aliquots were placed in cryovials for initial freezing at −80°C, after which they were transferred to liquid nitrogen for long-term storage.

### Cell culture

HEK-293T (ATCC, CRL-3216) were cultured according to ATCC recommendations in a humidified incubator maintained at 37°C and 5% CO2. HEK293T cells were cultured in DMEM supplemented with 10% FBS and 1% penicillin-streptomycin (Sigma-Aldrich, P0781), referred to as complete DMEM. T cells are cultured in X-VIVO 15 media (Lonza, 04-418Q) supplemented with 5% human serum (Sigma-Aldrich, H4522), 1% penicillin-streptomycin, 1% L-glutamine (Gibco, 25020-081), referred to as complete X-VIVO 15. IL-2 (Gibco, 200-02-100UG) was also added at 100 IU/mL. All FBS added into medium was heat-inactivated at 37°C for 30 min.

### Virus production

HEK293T cells were seeded 24 h before transfection at a density of 2 × 10^6^ cells per 100 mm Cell Culture Treated Dishes (Corning, 430167) in 10 mL Opti-MEM (Gibco, 31985-070) with 5% FBS, referred to as complete Opti-MEM. For each dish, plasmid DNA consisting of VR or CAR expression vectors, psPAX2 (Addgene #12260), and pMD2.G (Addgene #12259) was prepared at a ratio of 2 μg : 2 μg : 1 μg. DNA was diluted in 500 μL serum-free Opti-MEM and mixed with 15 μL GeneJuice Transfection Reagent (Merck Millipore, 70967). The mixture was incubated at room temperature for 5 min and added dropwise to the cells. Transfected HEK293T cells were maintained at 37 °C with 5% CO_2_, and the transfection mixture was replaced with fresh complete Opti-MEM after 8 h supplemented with 20 µL/10 mL ViralBoost Reagent (Alstem, VB100). Viral supernatants were collected at 72 h post-transfection, clarified by centrifugation at 500 × g for 10 min, and filtered through a 0.45 µm PES filter (Genesee, 25-246). Lentiviral particles were concentrated using the Lenti-X Concentrator (Takara, 631231). Viral supernatants were then mixed with Lenti-X Concentrator at a 3:1 ratio, incubated at 4 °C overnight, and centrifuged at 1,500 g for 45 min. The resulting pellet was resuspended in 1/100 of the original volume using serum free DMEM. Concentrated viral stocks were aliquoted and stored at –80 °C for future lentiviral transduction.

### T cell isolation and activation

Peripheral blood mononuclear cells were resuspended at 5 × 10^7^ cells/mL in EasySep™ Buffer (Stemcell, 20144) and transferred to a 5-mL polystyrene round-bottom tube (Falcon, 352054). The Isolation Cocktail (Stemcell, 17951C) was added at 50 µL/mL of sample, mixed, and incubated 5 min at room temperature. RapidSpheres™ (Stemcell, 50103) were added at 40 µL/mL of sample and mixed. The suspension was topped up to 2.5 mL with buffer and tube was placed into the EasySep™ magnet (Stemcell, 18000) and incubated 3 min at room temperature. The magnet and tube were then inverted to pour the enriched T-cell suspension into a new tube. Decant the enriched T-cell suspension into a new tube, centrifuged at 300 g for 5 min, and resuspend in X-VIVO 15 at 2 × 10^6^ cells/mL. Dynabeads Human T-Activator CD3/CD28(Gibco, 11131D) were added at 25 µL/mL and IL-2 at 100 IU/mL. Cells were seeded in 24-well plates (Falcon, 353047) at 1 mL/well and then incubated at 37°C with 5% CO_2_.

### Virus titration

Viral stocks were diluted 1:100 in serum free DMEM to a final volume of 150 µL. Viral RNA was isolated using the NucleoSpin RNA Virus Kit (Macherey-Nagel, 740956.50) according to the manufacturer’s instructions and eluted in 50 µL RNase-free water. Purified RNA was treated with DNase I to remove residual plasmid DNA, incubate at 37°C for 30 min, followed by 70°C for 5 min. Quantification of viral RNA copy number was performed using the Lenti-X qRT-PCR Titration Kit (Takara Bio, 631235) with standard curves generated from the provided RNA control template. Viral titers were calculated as RNA copies per milliliter following the manufacturer’s protocol.

### T cells transduction

96-well EIA/RIA plates (Corning, 3590) were pre-coated with 70 µL/well Retronectin reagent (Takara, T100A) at 40 µg/mL in DPBS (Gibco, 14190-144) and incubated at 4 °C overnight. Wells were washed with DPBS, lentivirus was added at the indicated volume, and each well was brought to 100 µL with serum-free DMEM. Plates were centrifuged at 2,000 g for 90 min at 32 °C. Viral supernatant was aspirated, and 1 × 10^5^ T cells were added per well in 100 µL viscous X-VIVO 15 containing 10% methylcellulose (R&D Systems, HSC001), 5% human serum, 1% penicillin– streptomycin, and 1% L-glutamine with IL-2 at 100 IU/mL and TransPlus Virus Transduction Enhancer at 2 µL/mL (Alstem, V050). After overnight incubation at 37 °C with 5% CO_2_, wells were washed 3 times with DPBS and replaced with 200 µL complete X-VIVO 15 containing IL-2 at 100 IU/mL. Transduction efficiency was assessed on days 6, 12, and 18 by flow cytometry, with medium refreshed the day before each measurement.

### Flowcytometry

Flow buffer was DPBS containing 0.5% w/v BSA (Sigma-Aldrich, A9647) and 0.05% w/v NaN_3_ (Sigma-Aldrich, S8032). 100 µL cell samples were washed 2 times in Flow buffer and resuspended to 120 µL. For fluorescent reporting cells, samples were filtered and acquired by flow cytometry. For antibody-stained cells, after one wash with Flow buffer, fluorophore-conjugated antibodies were added at 2 µL per antibody per well and incubated at 4 °C for 20 min protected from light. Cells were then washed 3 times with Flow buffer and resuspended in 120 µL Flow buffer. Flow cytometry was carried out on a Beckman Coulter CytoFLEX LX Flow Cytometer and data were analyzed with Flowjo Software version 10.10.

Antibodies used for labelling were as follows: APC AffiniPure F(ab’)_2_ Fragment Goat Anti-Human IgG, F(ab’)_2_ fragment specific (Jackson ImmunoResearch, 109-136-097), PE anti-human CD125 (Biolegend, 555902).

